# Distance estimation in the Goldfish (*Carassius auratus*)

**DOI:** 10.1101/2022.07.28.501828

**Authors:** Adelaide Sibeaux, Cecilia Karlsson, Cait Newport, Theresa Burt de Perera

**Affiliations:** Department of Biology, University of Oxford, Zoology Research and Administration Building, 11a Mansfield Road, Oxford, OX1 3SZ

**Keywords:** Distance estimation, Teleost, Optic flow, Navigation, Vision

## Abstract

Neurophysiological advances have given us exciting insights into the systems responsible for spatial mapping in mammals. However, we are still lacking information on the evolution of these systems and whether the underlying mechanisms identified are universal across phyla, or specific to the species studied. Here we address these questions by exploring whether a species that is evolutionarily distant from mammals can perform a task central to mammalian spatial mapping – distance estimation. We developed a behavioural paradigm allowing us to test whether goldfish (*Carassius auratus*) can estimate distance and explored the behavioural mechanisms that underpin this ability. Fish were trained to swim a set distance within a narrow tank covered with striped pattern. After changing the background pattern, we found that goldfish use the spatial frequency of their visual environment to estimate distance; doubling the spatial frequency of optic flow pattern resulted in a large overestimation of the swimming distance. These results provide robust evidence that goldfish can accurately estimate distance, and show that they use optic flow to do so. These results provide a compelling basis to utilise goldfish as a model system to interrogate the evolution of the mechanisms that underpin spatial cognition, from brain to behaviour.

## Introduction

Key neural structures that underpin navigation have been discovered in mammals, birds and reptiles (reviewed in Striedter, 2016). The ground-breaking discovery of place and grid cells in the late twentieth century changed the way we thought about the encoding of space (Hafting et al., 2005; O’Keefe & Dostrovsky, 1971; O’Keefe, 1976). These neural cells, located in the mammalian hippocampal formation, allow an individual to obtain information about its position in space by creating an internal map of its environment. Similar neural circuits have been found in birds (Atoji & Wild, 2006) and reptiles (Naumann et al., 2015) suggesting the existence of a common ground plan of a “hippocampal formation-like circuit” in ancestral amniotes (Striedter, 2016). Teleost fish are comprised of nearly 30 000 species (Ravi & Venkatesh, 2018), and are the most successful and diverse group of vertebrates, inhabiting various ecological niches and presenting a high level of morphological, physiological and behavioural diversity. However, the neurological structures and functions linked to spatial cognition in teleosts have only been investigated recently (see Rodríguez et al., 2021), and the neural basis of spatial cognition in teleost fish is still far from understood. This crucial information would allow us to build a more cohesive picture of the evolutionary origin of spatial navigation and its underpinning cells.

Neuroanatomical studies have shown that lesions in the lateral pallium affect the navigational performance of goldfish (*Carassius auratus*, Broglio et al., 2010; Rodríguez et al., 2002). Similar results were found when the hippocampal formation of rats was lesioned (Rodríguez et al., 2002), indicating that these structures are homologous. Neural cells that are likely to constitute the basic building blocks of fish navigation systems were recently discovered in the goldfish lateral pallium; head direction cells, edge detection neurons and speed correlated cells were recorded when individuals freely navigate in their environment (Vinepinsky et al., 2020). Crucially, we still do not know whether grid and place cells, the two main cell types used in mammalian spatial mapping and navigation, are present in teleost fish.

An essential step towards addressing this gap is to understand the spatial behaviour of teleost fish, the final output of spatial processing within the brain. Determining whether teleosts can navigate efficiently by computing moving distance and direction information, and the sensory mechanisms associated with these behaviours, will inform us about the neural structures that are likely to underpin them. Here, we use a behavioural paradigm to test whether the spatial metric of distance is encoded by a species of fish (Goldfish: *Carassius auratus*), with the intention of developing a new model system that can be used to link together behaviour with the neural mechanisms that drives it. Goldfish have already been used as a neural model for spatial cognition (Broglio et al., 2010; Durán et al., 2010; Rodríguez et al., 2021; Salas et al., 1996; Vinepinsky et al., 2020). They are easily accessible, and they are capable of learning, showing spatial, social and numerical cognitive abilities (Salena et al., 2021). in teleost fish. This study aims to first determine whether goldfish are able to encode distance travelled, making them a prime model system to explore space mapping in teleost fish and second, to test the sensory mechanisms behind this ability.

Previous results have shown that the Picasso Triggerfish (*Rhinecanthus aculeatus*) is able to accurately estimate travelled distance using frequency of the optic flow of their environment (Karlsson et al., 2019, 2020). Crucially, however, it is unknown whether goldfish, the potential model species to link neural and behavioural understanding of teleost navigation, possess the same abilities. It is possible that distance estimation is phylogenetically conserved across fish species. However, it is also possible that the environment might drive the evolution of the spatial mapping system. Goldfish and Picasso triggerfish inhabit very different environments and differ in many major behavioural traits. While the social goldfish inhabit murky water ponds, the territorial triggerfish are found in clear, bright reef water. The difference in how they might use space, and the visibility of navigational cues could lead to major divergence in their spatial mapping capabilities and underlying mechanism. We tested if (1) goldfish are able to accurately estimate travel distance, confirming their utility as a robust model species to investigate the neurophysiological basis of distance estimation in teleost fish; (2) they use the spatial frequency of optic flow to estimate distance, which we suggest as they are a highly visual species (Warburton, 1990); (3) alternative mechanism for distance estimation including travel time and number of fin beats.

## Material and Methods

### Experimental overview

We trained goldfish to a target distance in a long and narrow tank and then tested whether they could continue to swim the target distance given different optic flow information. In training, the fish were exposed to an achromatic striped pattern of 2cm and they were given an external cue to indicate when they reached the target distance. We then removed the external cue and measured if the fish continued to swim the set distance. Finally, we changed the optic flow pattern to determine whether the fish would change their estimate of distance travelled. Trial videos were then analysed to test whether alternative mechanisms, including fin beats and time, could have been used for distance estimation.

### Animal husbandry

Nine naive goldfish (*Carassius auratus*), sourced from a local supplier (The Goldfish Bowl, 118-122 Magdalen Rd, Cowley, Oxford OX4 1RQ) were used in experiments. Individuals were housed in 0.35m x 0.32m x 0.60m (width x height x length) tanks enriched with 0.5 cm of gravel, a terracotta pot, and plastic plants. Because *C. auratus* is a social species, individuals were kept in groups of two to three fish. The illumination by fluorescent light followed a 12h light /12h dark cycle. Individuals were fed twice a day; once in the morning with pellets (*Fancy Goldfish Sinking Pellets, FishScience*) and once in the afternoon with spinach or bloodworms to add supplementary nutrients. Tanks were cleaned weekly and water quality was maintained at healthy levels for this species (pH: 8.2; KH: 7dKH; GH: 8.2; Nitrite: 0ppm; Ammonia, Nitrate: <10ppm).

### Experimental apparatus

We used the experimental apparatus built by Karlsson et al. (2019). Briefly, fish were trained and tested in an acrylic tank (0.25m high x 0.16m wide x 1.80m length, Figure 1) set within a flow-through tank. The tank was connected to the home water system to maintain consistent water parameters, but the water flow was stopped during training and testing sessions. A white and black vertical 2 cm width stripe pattern (2 cm pattern) on the floor and walls of the tunnel produced optic flow cues. Three additional patterns were used to interrogate which optic flow features were used by the goldfish. 1) *Checker pattern*: a 2 cm^2^ checkerboard pattern with the same optic flow frequency as the training pattern was used to evaluate the impact of pattern change on fish distance estimation. 2) *High frequency optic flow pattern*: a 1cm width vertical stripe pattern, used to test the impact of increased optic flow on distance estimation. 3) *No optic flow pattern*: a 2cm horizontal stripe pattern used to test the impact of removing optic flow cues on distance estimation.

**Figure 1:**
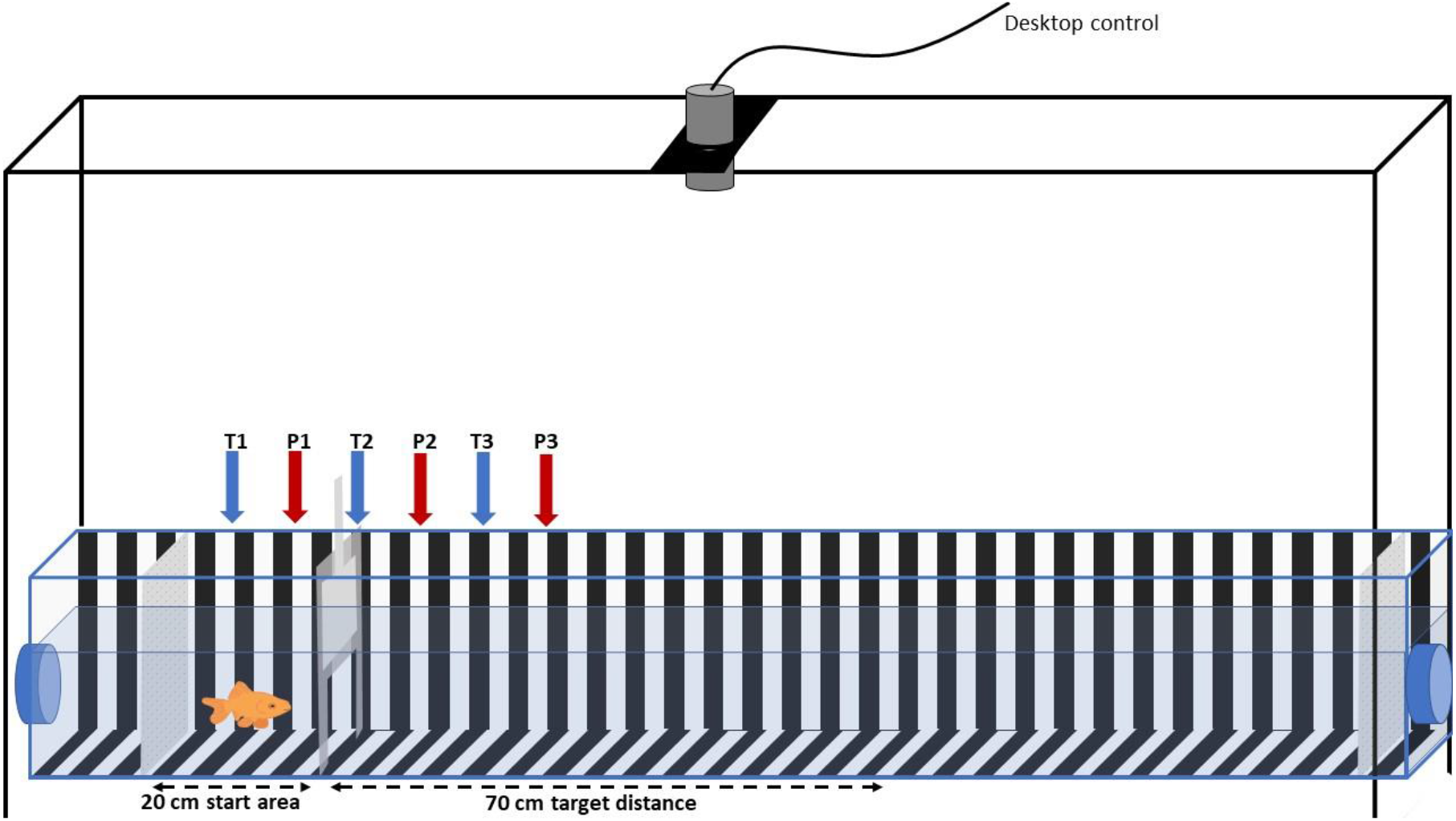
Experimental set-up to test distance estimation in the goldfish. Black and white striped panels (width 2 cm) covered the tank walls and floor providing constant optic flow information to the individual (only floor and left side wall are represented for clarity). The fish was placed in a movable start area, with a sliding door, for acclimation. Once the door opened the fish was trained to swim until the experimenter waved above the tank at the 70 cm target distance and then to return to the start area to receive a food reward. Two white partitions (grey on picture for clarity) prevented the fish to obtain external visual cues, one was placed at the end of the experimental tank and the other one was placed 20 cm behind the starting door. An overhead camera (grey cylinder) connected to the lab computer displayed fish movement in real time and recorded distance estimates during the testing phase. The linear tunnel was constructed inside a flow-through tank, with water flow in behind the start area (left bleu pipe) and passive water flows out at the opposite end of the tank (right blue pipe). Water flowed outside of training and testing sessions. T1, T2 and T3 indicate the three start positions used during training. P1, P2 and P3 indicate the three start positions used in test.

To control for the use of external visual cues to estimate distance, a moveable start area (0.25m high x 0.16m wide x 0.20m length) was placed at one of six start positions (10 cm apart). Three start positions (20 cm apart) were used to train the fish and three different start positions (20 cm apart) were used to test them. Therefore, while the target travel distance remained constant, the absolute position where the fish must turn was different in training and testing. A white partition was placed at the end of the tunnel to block external visual stimuli, and a second one was placed 20cm behind the start door. An overhead camera (Point Grey GrassHopper 3M-FLIR Machine Vision Cameras), placed 1.05m above the water level and connected to a computer displaying individual movement in real-time. All training and testing trials were recorded using StreamPix 7 video capture software (video frame rate = 50 fps).

### Training

We used an operant training paradigm with a food reward to train the fish to swim to a target distance of 0.70m. For each training session, the start area was randomly assigned to one of the three training start positions. During the first stage of training, bloodworms were placed along the bottom of the tank to encourage the fish to swim through. A transparent acrylic barrier was placed 70 cm apart from the door to stop the fish swimming further than the target distance. Bloodworms were then spaced throughout the tunnel to provide motivation for exploration. The number of bloodworms gradually decreased throughout training until only one blood worm was placed at the barrier and one at the start position. Gradually, the perplex barrier was replaced by a 5 cm partition, then a 0.5 cm stick, and finally the physical barrier was removed from the tank altogether. Those intermediate steps were necessary as they provided a physical cue indicating that the target distance was reached but also allowed the fish to get around it, swim further than the target distance and explore the tank. If the fish swam further than the target distance, no food reward was provided when it came back to the start position. The experimenter stood 1m apart from the experimental tank, observing the fish movements on the computer screen. The experimenter was not visible to the fish. The experimenter waved at the fish when it reached the target distance (with the 5 cm partition, 0.5 cm stick and without a physical barrier) to provide a cue for turning. At the final stage of training, the fish had to swim and turn when the experimenter waved above the experimental tank and then come back directly to the start position to receive their food reward. If the fish returned to the start position before reaching the target distance or explored the tunnel further away, no food reward was given. The training was completed once a fish reached the target distance in 80% of trials (i.e., swim out to the wave and come back directly to the start area) over three consecutive sessions. Training sessions lasted for 10 minutes or until 10 trials were completed. Fish were trained twice a day, once in the morning and once in the afternoon, five days a week.

### Testing

During testing, three start positions (different from the three training start positions) were randomly assigned and 15 trials per fish were completed at each start position. As a result, 45 distance estimation tests were recorded for each fish. For each session, fish first performed six to seven training trials with the experimenter providing a turning cue. The training start position was then moved to the test start position and the fish were tested for three to four trials with no turning cues. A food reward was given in all test trials regardless of the distance travelled. To test alternative optic flow patterns, fish were returned to their home tank after training while the experimenter changed the optic flow pattern. Fish were then moved back into the experimental tank and tested for a further three to four trials. A total of 45 distance estimation tests were also recorded for each optic flow pattern.

A few exceptions led to discarding a trial: (i) if the fish reached the very end of the experimental tank or if it only got its body length out of the start position before turning. (ii) If the fish turned multiple times in the tunnel before returning to the start position. (iii) If the fish showed erratic swimming movement indicating stress for the individual. In this final case, the trial was discarded, the session was ended and the fish returned to its home tank and monitored. Six of the nine fish were tested with the four different patterns. The power achieved to test the effect of the change of optic flow pattern on distance travelled with six individuals was above the 80% threshold criteria (power for 1000 iterations of the generalised linear mixed model= 94,50%, confident interval_95%_ [92.90; 95.83], R package *simr*, Green et al., 2016).

### Data collection

The videos were recorded using StreamPix7 software. To measure the goldfish distance estimate for each test trial, frames were extracted from video recordings at the point when the fish exited the start area and when it turned in the experimental tank. The pixel coordinates of the fish mouth were then manually recorded for those two events using custom video tracking software (Matlab version R2022a, MathWorks Inc.). The difference in pixel coordinate between the exit of the start area and the turn position was then converted into a distance estimate measured in centimetres (ratio 1 pixel = 0.0664 cm). The absolute turn position was measured using the coordinate of the turn frame converted in cm.

The time taken to turn in seconds was measured using the number of frames elapsed between the exit of the start area and the turn position. The frame rate (50 fps) was used to convert the number of frames to seconds. The number of caudal fin beats for each test trial was manually counted by the experimenter using StreamPix software that allowed frame-by-frame video inspection.

To determine if the goldfish used optic flow to estimate distance we compared the average turning distance and the absolute turn position obtained with the 2cm checker pattern (same density optic flow), the 1cm vertical stripes pattern (high frequency optic flow) and the horizontal stripes pattern (no optic flow).

### Statistical analyses

All statistical analyses were performed in *R* (version 4.0.2, 2020) with *R Studio* (R Studio 2009-2020, Version 1.3.1056). Normality and homogeneity of the residuals were successfully verified for each linear mixed model before further analysis. For each model detailed below (GLMM and LMM), we tested three different model structures with either “individual” as a random intercept, “individual” and “start position” as crossed random intercepts or “individual” as a random intercept and “start position” as a random slope. We compared models using the AIC criteria and performed analyses of variance (ANOVAs) between each model pair (Zuur et al., 2009) to select the best model structure. We also ran a post-hoc analysis on the selected model to control for multiple comparisons using the glht function and Holm-Bonferroni adjustment (R package *multcomp*, Hothorn et al., 2017). A power test (R package *simr*, Green et al., 2016) was performed post-hoc to verify that the number of individuals tested was sufficient to produce enough statistical power for following each changed in optic flow pattern. All tests were conducted with alpha = 0.05. We used the package *ggplot2* to draw figures (Wickham, 2016).

#### Overall distance estimation accuracy

To evaluate the goldfish distance estimation accuracy, individual average distance travelled and standard deviation was measured. We also performed a one-sample t-test on the 9 average values against the target distance mu=70. Normality of the data was preliminarily verified using a Shapiro-Wilk test (W=0.97, P=0.915).

As a control we tested if the order of the trial test had an effect on the distance travelled. A linear mixed-effects model (R package *lme4*, Bates et al., 2014) was performed with distance travelled included as the response variable and test order as the fixed effect. The model structure that fitted our data best included “individuals” and as random intercept.

#### Robustness of distance estimation across start position

To evaluate if the fish turned at the estimated target distance or if it learnt to turn using a landmark cue outside or inside the experimental tank, we tested if the start position affected the absolute turning distance. We performed a linear mixed-effects model (lmer package lme4, Bates et al., 2014) with absolute turning distance as the response variable and start position as the fixed effect. The model structure that fitted our data the best included “start position” and “individuals” as a random slope and intercept respectively.

#### Alternative cues used for distance estimation

To test the effect of three alternative cues: start position, number of fin beat and travel time on distance travelled, we performed a linear mixed model (lmer package lme4, Bates et al., 2014) with distance travelled as the response variable and start position, fin beats number and time as the fixed effects. The model structure that fitted our data the best included “individuals” and as a random intercept.

Because travel time and fin beat number are likely to increase with travelled distance, we measured the relative variation of distance travelled and either travel time or fin beat number. We measured the coefficient of variation (standard deviation/average x 100) of distance travelled vs. travel time/fin beats, and then determined the ratios.

As a control, we tested if the order of the test trial had an effect on the travel time. We performed a generalised linear mixed model (R package *lme4*, Bates et al., 2014) with a Gamma family to account for the elongated tail structure of the travel time data. Travelled time was included as the response variable and test order as the fixed effect. The model structure that fitted our data the best included “start position” and “individuals” and as a random slope and intercept respectively.

#### The use of optic flow information

To evaluate if goldfish use optic flow to estimate distance, we performed a linear mixed model (lmer package lme4, Bates et al., 2014) with distance travelled as the response variable and optic flow pattern as the fixed effect. The model structure that fitted our data the best included “start position” and “individuals” and crossed random intercept.

We ran a linear mixed model (lmer package lme4, Bates et al., 2014) to evaluate if the swimming speed (distance travelled/time) was affected by the optic flow patter. Speed was added in the model as the response variable and optic flow pattern as the fixed effect. The model structure that best fit our data included “individuals” as a random intercept.

## Results

### Overall distance estimation accuracy

All nine goldfish were able to reach the training criteria within 3 to 5 months (approximately 120 sessions and 1200 trials per fish) and complete testing. On average, the goldish swam 73.99 ± 16.77 cm (mean ± standard deviation, target distance = 70 cm, n= 405 trials) before turning (Figure 2, Table A1, Figure A1). Only 1 individual (Fish ID: G3), swam an average of 84.75 ± 13.92 cm and did not show an overlap of the target distance of 70 cm. The grouped mean distance travelled was not significantly different from that of the target distance (one-sample t-test, t=2.12, df=8, p=0.066, CI_95%_ [69.65 ; 78.32]). We did not find any effect of the test order on distance travelled (P=0.494, see Table A2 for details).

**Figure 2:**
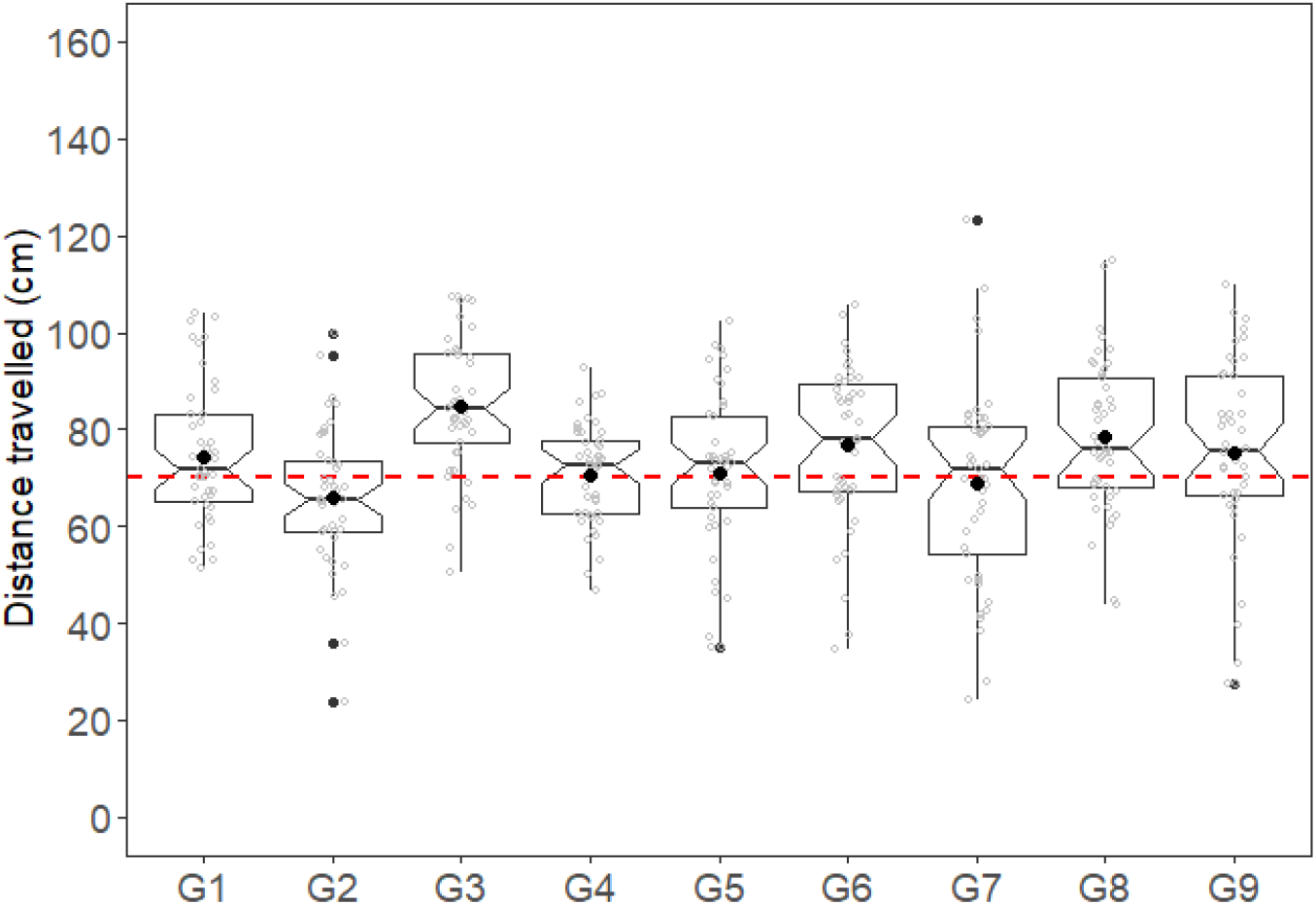
Distance estimated for the nine goldfish tested with the 2cm optic flow pattern. Black dots represent the mean distance estimated for each fish. The red dashed line at 70 cm represents the target distance. Raw data are represented by the grey dots. See Table A1 and Figure A1 for details. Histograms of the distance estimate are presented in Figure A2.

### Distance estimation is robust to changes in start position

The start positions significantly affected the individual absolute turn distance (turning point in the experimental tank) (Figure 3a, Table 1). As predicted if fish were accurately estimating distance, the further the fish started within the tank, the further they turned. Absolute swimming distances were 104.86 ± 18.25 cm, 120.93 ± 16.55 cm and 134.06 ± 13.40 cm when the fish started at the 1^st^, 2^nd^ and 3^rd^ position respectively.

**Figure 3:**
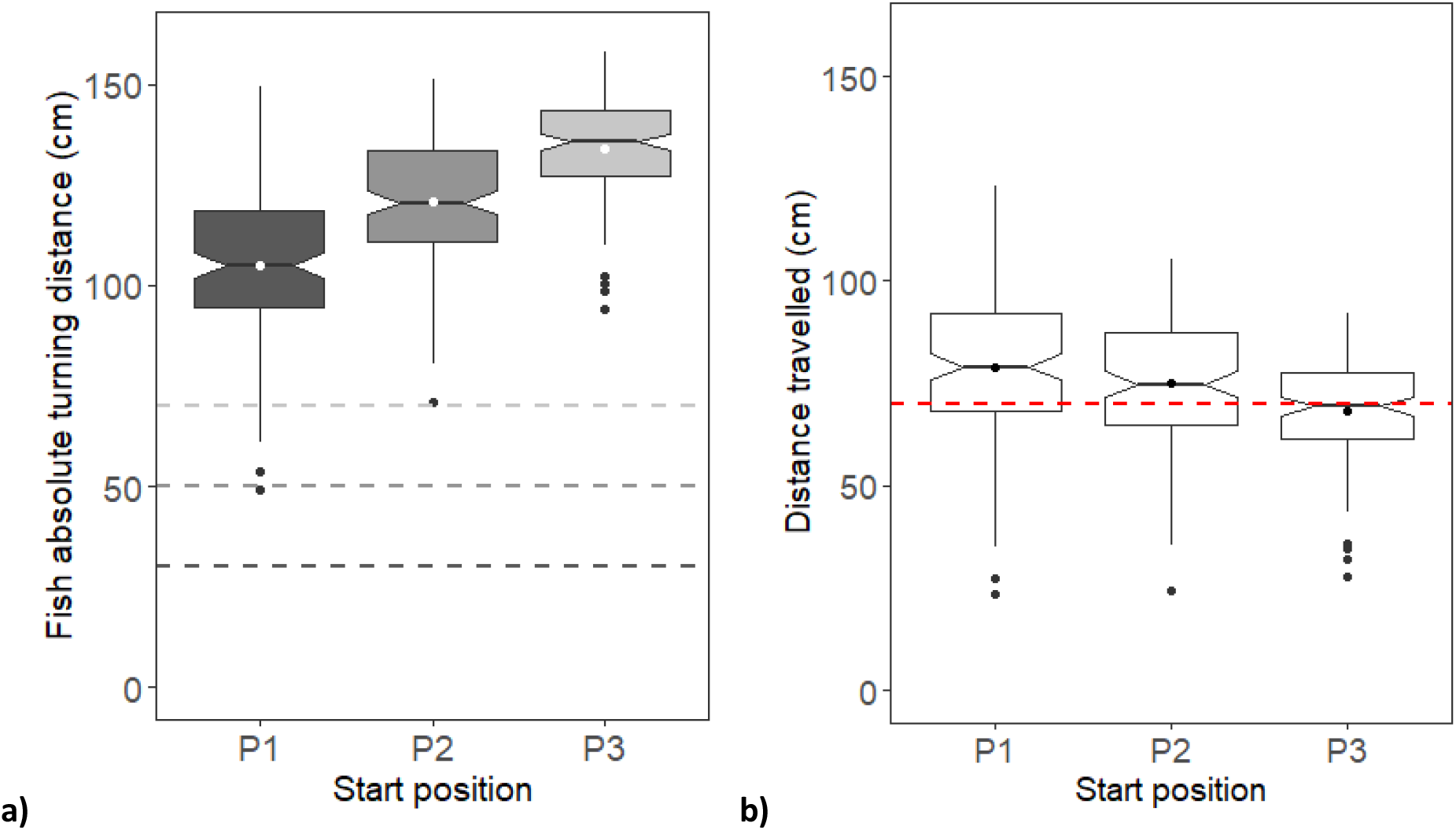
Goldfish distance estimation accuracy as a function of start distance with a 2 cm optic flow pattern. a) Absolute turning distance at three start positions. The dashed lines represent the three start positions. White dots represent the mean absolute turning distance for each start position. b) Distance travelled per start position. The black dots represent the mean distance travelled. The red dashed line represents the 70 cm target distance. See figure A3 for individual details.

**Table 1:** Effect of the start position on the absolute turn position in the experimental tank. Adjusted p-values (Holm method) are reported.

| Fixed effects | $\beta$ coefficient | SE | df | t | P |
| --- | --- | --- | --- | --- | --- |
| Intercept | 104.86 | 2.87 | 8.00 | 36.59 | **<0.001** |
| Position(P2-P1) | 16.07 | 2.75 | 8.07 | 5.85 | **<0.001** |
| Position (P3-P1) | 29.20 | 3.07 | 8.09 | 9.53 | **<0.001** |
| Position (P3-P2) | 13.13 | 2.53 | 9.94 | 5.19 | **<0.001** |
| Random effects | Variance | SD | R |  |  |
| Individual (intercept) | 58.85 | 7.67 |  |  |  |
| Position(P2-P1) | 37.77 | 6.15 | -0.43 |  |  |
| Position (P3-P1) | 54.40 | 7.38 | -0.94 |  |  |
| Position (P3-P2) | 27.47 | 5.24 | -0.98 |  |  |
| Residuals | 225.95 | 15.03 |  |  |  |

Results from the linear mixed model with fish ID as random intercept and position as random slope. Significant results are in bold. df = degrees of freedom; $P$ = $P$ value; SD = standard deviation; SE = standard error; $t$ = $t$ value.

### Alternative cues used for distance estimation

We tested the effect of start position, fin beats number and time on goldfish distance travelled. Fish swam a significantly shorter distance when they started from the 3^rd^ position (68.06 ± 13.43 cm) than from the 1^st^ (78.82 ± 18.24 cm) or 2^nd^ (75.07 ± 16.56) position (Table2, Figure 3b). There was no significant difference between the 1^st^ and 2^nd^ start position. Individual were closer to the target distance and more accurate in their distance estimate at the 3^rd^ position. Individual distances travelled at each start position are presented in Appendix Figure A3.

**Table 2:**
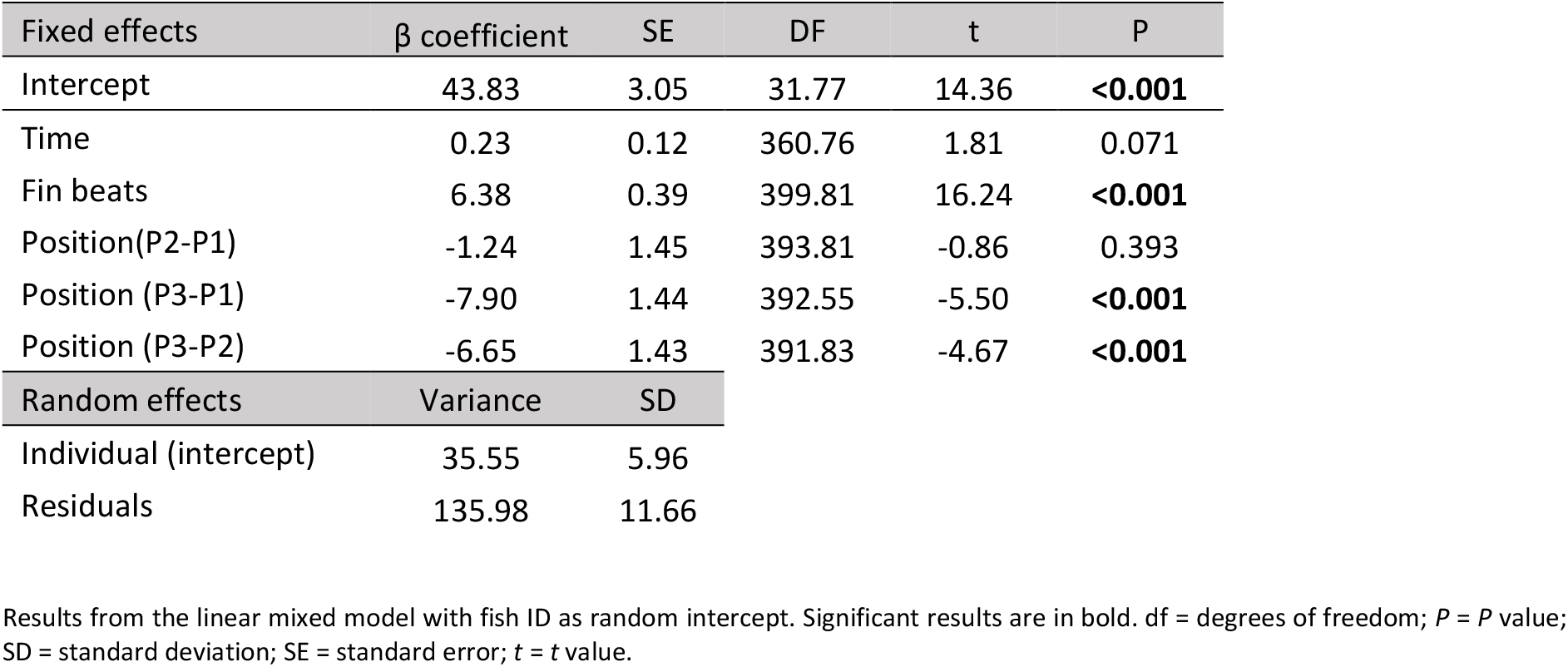
Effect of travel time, number of fin beats and start position on goldfish distance travelled. Adjusted p-values (Holm method) are reported.

The number of caudal fins beats significantly and positively correlated to the swimming distance (Table 2, Figure 4), so that the greater the distance travelled, the more fin beats occurred. The ratios of the coefficient of variation between fin beats number and distance travel are ranging from 1.12 and 1.94. Ratios close to one indicate that the fish were similarly consistent in their distance estimation and their fin beats number across trials. Ratios closer to two indicate that the variation in the number of fin beats produced was twice as large as the variation in the distance estimate (Table 3).

**Figure 4:**
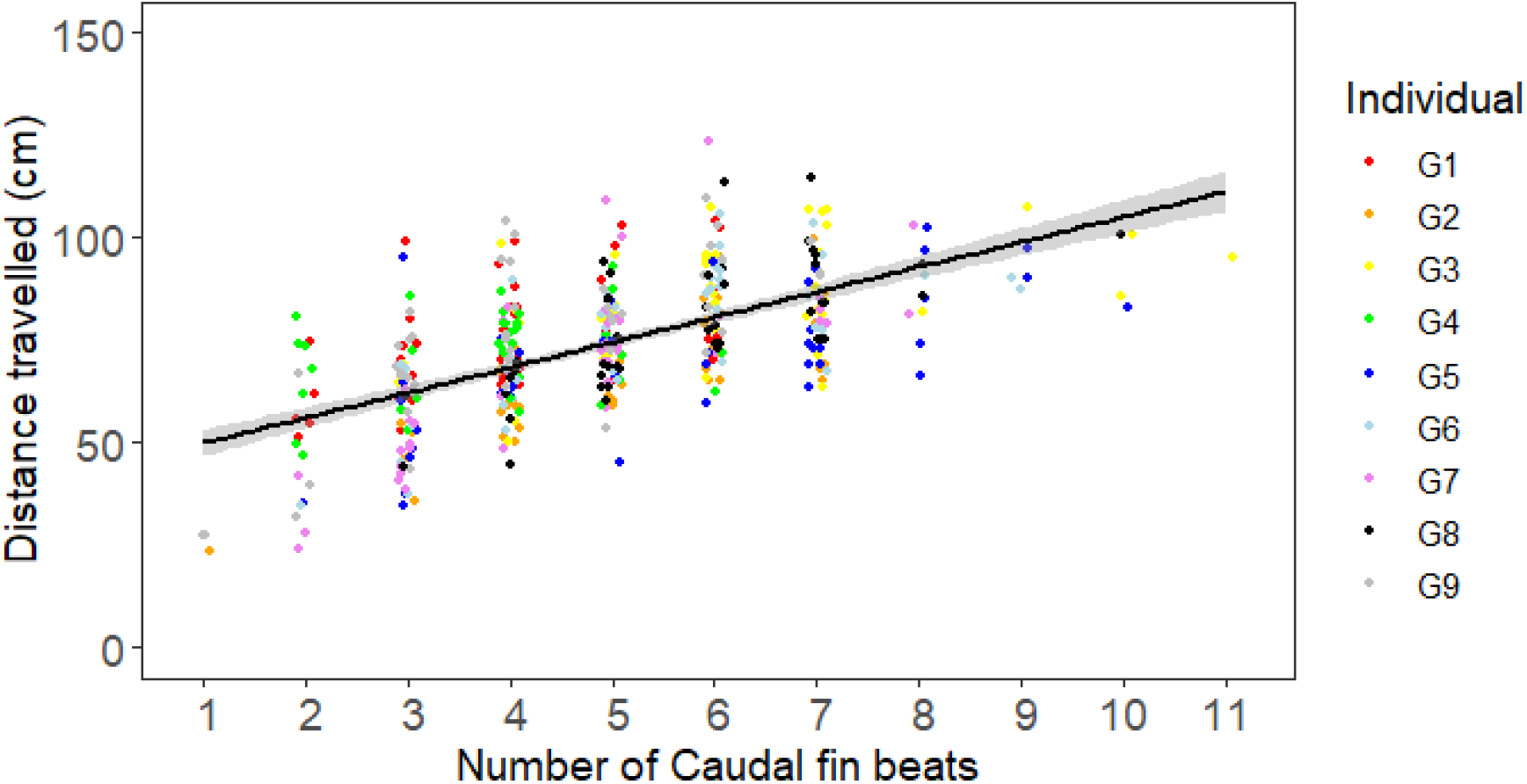
Effect of the number of caudal fin beats on the travelled distance. The black line and grey shading represent the linear regression ± SE. The positive slope R=62.3% indicates that fin beats is positively correlated to distance travelled.

**Table 3:** Coefficient of variation of distance travelled and number of fin beats for each individual.

| Individual | Distances travelled CV | Number of fin beats CV | CV Ratio |
| --- | --- | --- | --- |
| G1 | 19.51 | 30.08 | 1.54 |
| G2 | 21.91 | 28.74 | 1.31 |
| G3 | 16.42 | 28.64 | 1.74 |
| G4 | 14.57 | 28.21 | 1.94 |
| G5 | 23.16 | 35.75 | 1.54 |
| G6 | 21.34 | 30.52 | 1.43 |
| G7 | 29.94 | 33.67 | 1.12 |
| G8 | 20.04 | 24.50 | 1.22 |
| G9 | 26.35 | 37.25 | 1.41 |

We did not observe a significant effect of travel time on goldfish distance travelled (Table 2). Moreover, the ratio of coefficient of variation between the travel time and the distance travelled ranged from 1.29 to 3.03 (Table 4). On average, the ratio of coefficient of variation was twice as large for travel time than for distance travelled indicating that goldfish were much more consistent in their travel distance estimation than in their travel time. We did not find any effect of the test order on the travel time of individuals (P=0.614, see Table A3 for details).

**Table 4:** Coefficient of variation of distance travelled and travel time for each individual.

| Individual | Distances travelled CV | Time CV | CV Ratio |
| --- | --- | --- | --- |
| G1 | 19.51 | 37.51 | 1.92 |
| G2 | 21.91 | 31.15 | 1.42 |
| G3 | 16.42 | 31.25 | 1.90 |
| G4 | 14.57 | 22.95 | 1.57 |
| G5 | 23.16 | 58.58 | 2.53 |
| G6 | 21.34 | 49.50 | 2.32 |
| G7 | 29.94 | 43.77 | 1.46 |
| G8 | 20.04 | 25.89 | 1.29 |
| G9 | 26.35 | 79.78 | 3.03 |

### The use of Optic Flow Information

Six individuals were tested with the four previously described optic flow patterns: 2 cm, checker, high frequency, and no optic flow. We did not find a significant difference in distance travelled when goldfish were tested with the 2 cm (73.94 ± 15.51 cm) or the checker pattern (74.94 ± 18.79 cm, Figure 5, Table 5). The fish swam a significantly shorter distance (47.46 ± 21.47 cm) when they were tested with the high frequency optic flow pattern than with any other pattern. The distance travelled with the no optic flow pattern (65.04 ± 30.55 cm) was significantly shorter than with the 2 cm and checker patterns and significantly longer than with the high frequency optic flow pattern. Moreover, the goldfish showed much more variability in their distance estimate with no optic flow pattern, with a standard deviation twice as large as for the 2 cm pattern. Individual travel distance for each pattern are given in the Appendix Table A1. Details of the distance travelled with the four different patterns for each of the six individual and for the three start positions are presented in Figure A4 and A5 respectively. Goldfish swimming speed was also significantly slower when they were tested with the high frequency (7.44 ± 3.21 cm/s) and no optic flow (5.95 ± 2.57 cm/s) patterns, compared to the 2cm (11.08 ± 4.75 cm/s) and checker patterns (11.28 ± 3.99 cm/s, Figure 6, see Table A4 for p-values and model summary).

**Figure 5:**
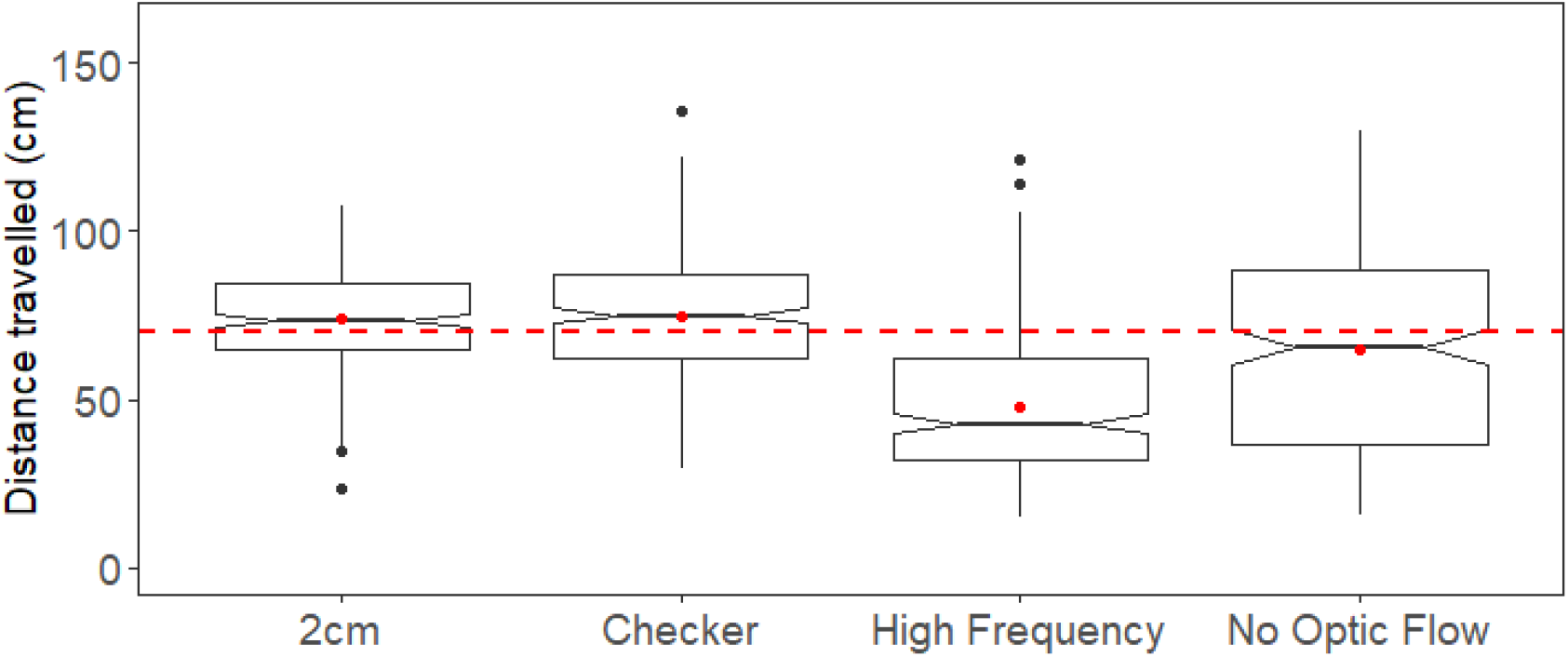
Summary of estimated distance for the four optic flow patterns. Red dashed line represents the 70cm target distance. Red dots indicate the mean distance estimate for each pattern. N=6 fish, see Figure A4 for individual details

**Table 5:** Effect of the optic flow pattern on the travel distance.

| Fixed effects | $\beta$ coefficient | SE | DF | z | P |
| --- | --- | --- | --- | --- | --- |
| Intercept | 73.94 | 4.16 | 7.69 | 17.77 | **<0.001** |
| Checker - 2cm | 1.00 | 1.79 | 1069 | 0.56 | 0.576 |
| High Frequency - 2cm | -26.48 | 1.79 | 1069 | -14.81 | **<0.001** |
| No Optic Flow - 2cm | -8.90 | 1.79 | 1069 | -4.98 | **<0.001** |
| High Frequency - Checker | -27.48 | 1.79 | 1069 | -15.36 | **<0.001** |
| No Optic Flow - Checker | -9.90 | 1.79 | 1069 | -5.53 | **<0.001** |
| No Optic Flow - High Frequency | 17.58 | 1.79 | 1069 | 9.83 | **<0.001** |
| Random effects | Variance | SD |  |  |  |
| Individual (intercept) | 66.89 | 8.18 |  |  |  |
| Position (intercept) | 13.67 | 3.70 |  |  |  |
| Residuals | 431.85 | 20.78 |  |  |  |

Results from the linear mixed model with fish ID and position as crossed random intercept. Significant results are in bold. df = degrees of freedom; P = P value; SD = standard deviation; SE = standard error; t = t value.

**Figure 6:**
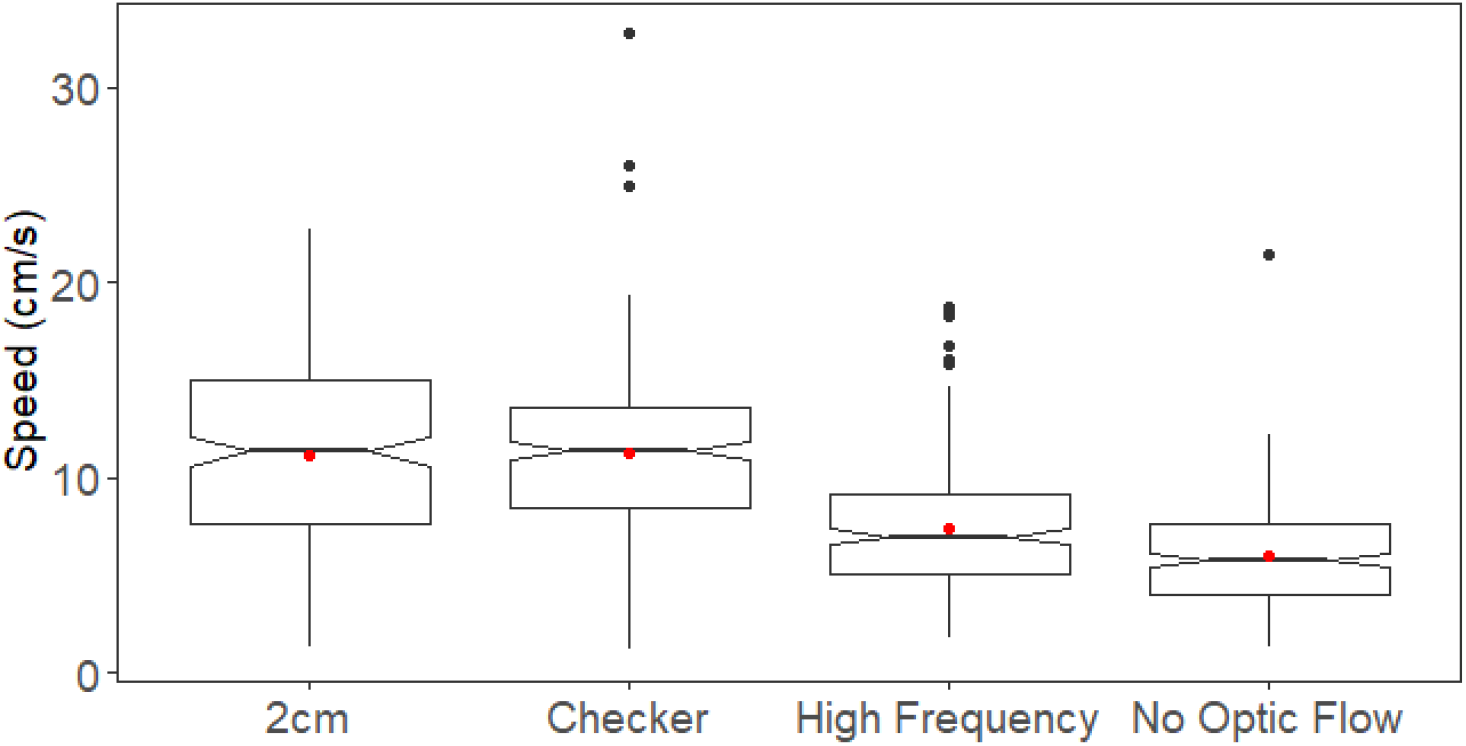
Goldfish swimming speed across optic flow treatments. Red dots indicate the mean distance estimate for each pattern. Fish tested n=6.

## Discussion

Distance is an important metric that underpins spatial cognition, and we suggest that this behaviour can be exploited to study the evolution of spatial cognition from behaviour through to the neural systems that underly it. In this study, we have found that goldfish are able to accurately reproduce a learned distance and that they can use optic flow cues to do so. Goldfish were trained to travel a distance of 70 cm before returning to a start position and continued to do so when external cues (experimenter waving) were removed (average swimming distance =74.0 ± 16.7 cm). Although distance estimation accuracy was impacted by start position, it is unlikely that fish used a fixed landmark cue (external or internal to the experimental tank) for distance estimation because they were able to swim the approximate learned distance at all start positions (20 cm apart) and therefore were turning at different absolute positions depending on the start position. This first result indicates that the spatial metric of distance is encoded by goldfish and that this species provides a robust model system for future examinations of the neural basis of spatial cognition in teleost fish, allowing to link together behaviour with the neural mechanism that drives it.

The ability to estimate distance in fish was recently shown in Picasso Triggerfish (Karlsson et al., 2019), but there were some interesting differences between that species fish and the goldfish. Firstly, the goldfish were not as accurate as the triggerfish when estimating distance travelled. The triggerfish were trained to swim 80 cm and travelled an average of 80.3 ± 3.7 cm. Multiple factors could explain this difference. Firstly, the triggerfish may encode distance more accurately than goldfish. This difference in ability could simply be species-specific but could also be related to the difference in environmental rearing conditions (see Ebbesson & Braithwaite, 2012 for a review). While the goldfish were reared in captivity, the triggerfish came from a wild population and therefore were exposed to a higher variety of environmental stimuli. It has been shown that rearing and enrichment conditions can significantly impact forebrain development and cognitive abilities in salmon (Kihslinger et al., 2006; Salvanes et al., 2013). In Chinook Salmon (*Oncorhynchus tshawytscha*), wild-caught individuals had significantly larger olfactory bulb and telencephalon volumes relative to their body size compared to individuals reared in hatcheries (Kihslinger et al., 2006). In Atlantic Salmon (*Salmo salar*), individuals reared in enriched environments (e.g., pebbles, rocks and floating structures) were more accurate at locating the exit of a maze compared to individuals reared without (Salvanes et al., 2013). Secondly, triggerfish might be more accurate in distance estimation because of the ecological relevance of the task. This territorial species requires accurate spatial memory to travel back to their own home after foraging trips. Moreover, knowledge of the position and distance to the nearest shelter would be valuable when encountering predators. Finally, small differences in experimental training and testing could have had an effect. For example, when the triggerfish reached the target distance, it triggered an infrared detector and activated a flashing light training cue. This training set up was not possible for the goldfish as the detector was not accurate for their given size, and propensity to swim at the bottom and out of range of the infrared signal. Instead, the experimenter waved at the goldfish to indicate that the target distance was reached. The millisecond of latency for the experimenter to wave might have given the fish a less accurate idea of the target distance.

The modification of the spatial frequency of the background pattern significantly affected the distance travelled by the goldfish, providing evidence for the use of optic flow as a cue for distance estimation. When the optic flow pattern frequency increased (high frequency optic flow pattern), goldfish overestimated the distance they travelled by 36%, and turned before reaching the target distance. Goldfish demonstrated the same behavioural pattern when presented with the checker pattern (same optic flow frequency than the 2 cm pattern) and the 2 cm pattern, indicating that it was the change of optic flow frequency and not the change of pattern that altered individual swimming distance. In the absence of optic flow information (no optic flow pattern), the fish distance estimate became inconsistent, with a standard deviation that doubled compared to the 2 cm optic flow pattern. These results suggest that the visual odometer of goldfish is mediated by a movement detection mechanism similar to that underlying optomotor response (Arnold, 1974). The use of optic flow to estimate distance travelled is widespread in other vertebrate and invertebrate species. Humans, ants, wolf-spiders and honeybees also use optic flow as a visually guided odometer (Ortega-Escobar & Ruiz, 2014; Redlick et al., 2001; Ronacher & Wehner, 1995; Srinivasan et al., 1997). However, they integrate the image motion (i.e., speed of visual background on the retina) and not the structure of the background (i.e., frequency of the optic flow) to estimate distance. Those species therefore used “true optic flow” and measured the angular speed of visual features across the retina to estimate travelled distance. In those cases, the change in spatial frequency did not affect individual distance estimation scores. However, changes in either speed or distance to the visual background would affect distance perception. These results combined with our own, indicate that visually-based distance estimation is widely spread across taxa but that the visual information extracted (frequency versus angular speed) differs from fish compared to insect or mammals. The similarity of goldfish and triggerfish behaviour in response to changes in optic flow cues (i.e. high frequency pattern = overestimated travel distance, no optic flow = reduced accuracy) Karlsson et al., 2020) suggests that the use of background spatial frequency information as an optometer may be widespread amongst teleosts. Particularly given that the two species inhabit different visual environments (Cheney et al., 2013; Mitchell et al., 2017; Newport et al., 2017).

In contrast to the results obtained with triggerfish, goldfish swimming speed was significantly affected by changes in the spatial frequency of the optic flow pattern. In goldfish, the travel distance and travel speed both decreased by 36% and 33% respectively when the optic flow frequency was doubled. We could wonder if goldfish were using time as a proxy to estimate distance; swimming a shorter distance slower. However, supplementary analyses showed that goldfish swimming time was significantly different for each optic flow pattern tested. If individuals were using time as a proxy to estimate distance, we would expect the travel time to be consistent across the optic flow pattern tested. Instead, we found that individuals swam a significantly longer amount of time with the 2cm optic flow pattern (Table A5 for details). Because travel distance and travel speed were both affected by the optic flow frequency we can assume that the rate of movement of visual contrast might be used in the same way for speed control and for odometry. The visual control of swimming speed and odometry by goldfish could therefore be underpinned by the same visual motion mechanisms. This mechanism could be shared with other species such as the honeybees, who use optic flow to control their speed (Portelli et al., 2011). The use of optic flow information allows honeybees to adjust both their speed and distance travelled when foraging in complex environments and a similar mechanism may confer similar benefits to fish. Interestingly, for teleost fish speed correlated cells have recently been found in the goldfish lateral pallium when they are freely navigating within an a confined space (Vinepinsky et al., 2020). However, there was no optic flow pattern present in this study so the lateral line was the most likely sensory system to inform individuals about their swimming speed. Our results indicate that the information gathered by both the lateral line and the individual visual system could inform goldfish about their swimming speed. A recent review pointed out that spatial memory in teleost fish could be possibly considered a special case of a wider relational memory system that encodes both the spatial and the temporal dimensions of episodic-like memories and that those encoding take place in the hippocampal pallium of teleost fish (Rodríguez et al., 2021).

Along with optic flow information, goldfish could use alternative cues to estimate distance. We explored the role of the start position, the number of fin beats and the travel time in goldfish distance estimation. Goldfish performed significantly better (mean travelled distance closer to the target distance and smaller standard deviation) when tested at the third position than at the first or second position. When tested at the third position, the absolute turn position was closer to the end of the experimental tank making it impossible to swim beyond 30 cm after the 70 cm target distance before reaching the end of the experimental tank. When tested at the first start position they were able to swim 70 cm after the 70cm target distance. This might be a confounding effect leading to a better accuracy (lower standard deviation) when tested at the third position. Moreover, it is possible that individuals used a combination of self-integrated distance measurement and spatial cues to obtain better accuracy in their travelled distance estimate. The closeness to the end of the tunnel could give them supplementary information about their turning point. Studies have shown that multiple fish species (e.g., Redtail Splitfins *Xenotoca eiseni*, and goldfish) were able to use the geometry of space (geometrical cues of a rectangle arena) in navigation and decision making (Sovrano et al., 2002, 2003; Vargas et al., 2004). In a spontaneous reorientation task (i.e., where no reinforced training was used) Redtail Splitfins and Zebrafish (*Danio rerio*) could also use geometrical information to find the previous position of a conspecific (Lee et al., 2012; Sovrano et al., 2020). Recent neurophysiological work by Vinepinsky et al. (2020) has revealed the presence of edge detection neurones in the lateral pallium of goldfish, which is a first indicator that teleost fish are able to encode information about the features of space.

The number of caudal fin beats was significantly correlated to the swimming distance. The ratio of the variation of fin beat number and travelled distance ranged from 1.1 to 1.9. Variation in fin beat number was therefore greater than the variation in travelled distance for each individual. However, for 5 individual the ratio was below 1.5. Individuals with a ratio of 1.1 and 1.2 had similar variation in distance estimate and fin beat number. It is thus possible that some individuals used fin beats as an odometer and a proxy to estimate travel distance. Individual differences in navigation strategies have been observed in other fish species. For example, Lee et al. (2012) found species (Redtail Splitfins versus Zebrafish) and sex difference in the use of landmark in a reorientation task. While both sexes used geometrical cues to orient, the combined use of landmark and geometry was only significant in males. Females only showed the use of geometry without distinction between the two geometric twins. Males using both geometrical and landmark cues were therefore more accurate but only when the goal was nearer to the attractive local landmark (they used the landmark as a beacon). Moreover, Redtail Splitfins were successful at using both landmark and geometrical cues while Zebrafish used mainly geometrical cues. The ratio of variation in fin beat number versus variation in distance travelled was much lower in goldfish than in the triggerfish (Karlsson et al., 2019). Species differences in navigational strategies could explain the fact that triggerfish were not using fin beats to measure distance. This species-specific strategy could be explained by the subcarangiform swimming pattern in goldfish and the oscillatory swimming style of triggerfish. In other species, while the honeybees (*Apis mellifera*) odometer is driven by image motion (Srinivasan et al., 2000), desert ants (*Cataglyphis)* and Fiddler Crab (*Uca pugilator*) use step count (pedometer) as an odometer (Walls & Layne, 2009; Wittlinger et al., 2006). A definitive answer as to whether goldfish use fin beats as odometer requires further investigation.

Goldfish were unlikely to use time as a proxy to estimate distance travelled. We did not find a significant effect of time on the distance travelled and the coefficient of variation in time travelled was two to three times higher than the coefficient of variation in distance estimate for most fish. Video analysis revealed high variation in time spent to perform the experiment within a day or a session without any apparent pattern, further supporting this hypothesis. The trial order did not significantly impact the time spent to perform the task. This result is consistent with triggerfish (Karlsson et al., 2019) that also did not use time as a proxy to estimate the travelled distance. Studies with other taxa also reported that time was uncorrelated with travelled distance. For example, honeybees did not use time to estimate flight distance when tested in a tunnel with either headwind or tail wind (Srinivasan et al., 1996; Srinivasan et al., 1997). Moreover, in humans, the variability at reproducing a given distance by walking was found to be lower than the variable to estimate an interval of time. This result did not favour a temporal metric for locomotor distance perception in humans because their precision in distance production was greater than their precision of temporal interval (Durgin et al., 2009). If individuals used time as a proxy to measure distance travelled, the error produced in the distance estimation task (variation in distance estimate) should be similar to the error in time to perform the task. However, in this experiment, the variability in distance travelled was two times lower than that of travel time. In their natural environment, time may not be an accurate proxy for distance as goldfish are likely to forage and interact with conspecific while travelling from a point A to a point B and therefore high temporal variability during their travel is likely to occur. Flow within aquatic systems could also disrupt this estimate.

In conclusion, we have shown that goldfish use the spatial frequency of the visual background for odometry and speed control. The two processes might be underpinned by a similar motion detection mechanism and are likely to share a similar neural pathway. This study identifies the goldfish as a robust model species to examine the neural basis of spatial cognition in teleost fish, and advances the use of this animal system to understand the evolution of spatial mechanisms (neural and behavioural).

## Acknowledgements

This research was funded thanks to the human frontier science program grant number RGP0016/2019. CN was funded by the Leverhulme Trust Early Career Fellowship. We thank Christine Soper for her help with animal husbandry.

## Ethical note

This experiment was conducted under Oxford University’s AWERB (Animal Welfare and Ethical Review Board) approval number APA/1/5/ZOO/NASPA. All individuals were handled with the highest care to minimize stress.

## Notes

### Competing Interest Statement

The authors have declared no competing interest.

