## Appendix for "Distance estimation in the Goldfish (*Carassius auratus*)"

### Supplementary material

Adelaide Sibeaux[1], Cecilia Karlsson[1], Cait Newport[1], Theresa Burt de Perera[1]

[1] Department of Biology, University of Oxford, Zoology Research and Administration Building, 11a Mansfield Road, Oxford, OX1 3SZ.

### Appendix

Table A1: Fish distance estimate average and SD

| Fish ID | 2cm | | checker pattern | | HFOF pattern | | NoOF pattern | |
|---|---|---|---|---|---|---|---|---|
| | Average Distance | Distance SD | Average Distance | Distance SD | Average Distance | Distance SD | Average Distance | Distance SD |
| G1 | 74.40 | 14.51 | 76.30 | 15.32 | 39.72 | 15.71 | 67.74 | 28.10 |
| G2 | 66.01 | 14.47 | 71.58 | 19.37 | 33.84 | 14.21 | 52.17 | 24.44 |
| G3 | 84.75 | 13.92 | 82.12 | 17.69 | 60.79 | 28.96 | 79.84 | 31.57 |
| G4 | 70.69 | 10.30 | 70.87 | 14.52 | 50.54 | 14.87 | 49.65 | 22.37 |
| G5 | 71.05 | 16.46 | 81.20 | 19.08 | 59.23 | 17.86 | 84.40 | 27.08 |
| G6 | 76.73 | 16.37 | 67.55 | 21.93 | 40.61 | 18.95 | 56.43 | 31.42 |
| G7* | 68.80 | 20.60 | | | | | | |
| G8* | 78.44 | 15.72 | | | | | | |
| G9* | 75.02 | 19.77 | | | | | | |
| Average all fish | 73.99 | 16.77 | 74.94 | 18.79 | 47.46 | 21.47 | 65.04 | 30.55 |

*G7, G8 and G9 were tested only with the 2cm optic flow pattern

Table A2: Effect of the test order on the distance travelled

| Fixed effects | β coefficient | SE | df | t | P |
|---|---|---|---|---|---|
| Intercept | 75.00 | 2.39 | 20.80 | 31.40 | **<0.001** |
| Test order | -0.04 | 0.06 | 396.16 | -0.69 | 0.494 |
| **Random effect** | **Variance** | **SD** | | | |
| Individual (intercept) | 25.95 | 5.094 | | | |
| Residuals | 258.25 | 16.07 | | | |

Results from the linear mixed model with fish ID as random intercept. Model selection on the random effect was conducted with ANOVAs (REML fitted) on pairs of models containing either “individual” as random intercept, “individual” and “start position” as crossed random intercept or “individual” as random intercept and “start position” as random slope. We used AIC criteria to select the best model .Significant results are in bold. df = degrees of freedom; *P* = *P* value; SD = standard deviation; SE = standard error; *t* = *t* value.

Table A3: Effect of the test order on the time to perform the distance estimation task

| Fixed effects | β coefficient | SE | t | P |
|---|---|---|---|---|
| Intercept | 0.13 | 0.02 | 8.60 | **<0.001** |
| Test order | $7.49 \times 10^{-5}$ | $1.49 \times 10^{-4}$ | -0.50 | 0.614 |
| **Random effects** | **Variance** | **SD** | **R** | |
| Individual (intercept) | $4.95 \times 10^{-4}$ | 0.02 | | |
| Position(P2-P1) | 0.03 | -0.29 | -0.29 | |
| Position (P3-P1) | 0.03 | 0.02 | 0.02 | |
| Position (P3-P2) | $2.21 \times 10^{-4}$ | 0.01 | 0.39 | |
| Residuals | 0.17 | 0.41 | | |

Results from the generalised linear mixed model, fitted with Gamma family. Fish ID was added as random intercept and start position as random slope. Model selection on the random effect was conducted with ANOVAs (REML fitted) on pairs of models containing either “individual” as random intercept, “individual” and “start position” as crossed random intercept or “individual” as random intercept and “start position” as random slope. We used AIC criteria to select the best model .Significant results are in bold. *P* = *P* value; SD = standard deviation; SE = standard error; *t* = *t* value.

Table A4: Effect of optic flow pattern on the swimming speed of goldfish.

| Fixed effects | β coefficient | SE | z | P |
|---|---|---|---|---|
| Intercept | 11.08 | 0.77 | 14.49 | **<0.001** |
| checker - 2cm | 0.20 | 0.29 | 0.69 | 0.489 |
| High Frequency - 2cm | -3.64 | 0.29 | -12.67 | **<0.001** |
| No Optic Flow - 2cm | -5.13 | 0.29 | -17.86 | **<0.001** |
| High Frequency - Checker | -3.84 | 0.29 | -13.36 | **<0.001** |
| No Optic Flow - Checker | -5.33 | 0.29 | -18.55 | **<0.001** |
| No Optic Flow - High Frequency | -1.49 | 0.29 | -5.19 | **<0.001** |
| **Random effects** | **Variance** | **SD** | | |
| Individual (intercept) | 3.27 | 1.81 | | |
| Residuals | 11.14 | 3.34 | | |

Results from the linear mixed model with fish ID as random intercept. Model selection on the random effect was conducted with ANOVAs (REML fitted) on pairs of models containing either “individual” as random intercept, “individual” and “start position” as crossed random intercept or “individual” as random intercept and “start position” as random slope. We used AIC criteria to select the best model .Significant results are in bold, *P* = *P* value; SD = standard deviation; SE = standard error; z = *z* value.

Table A5: Effect of optic flow pattern on swimming time of the goldfish

| Fixed effects | β coefficient | SE | z | P |
|---|---|---|---|---|
| Intercept | 0.12 | 0.01 | 9.20 | **<0.001** |
| checker - 2cm | 0.02 | $4.49 \times 10^{-3}$ | 3.74 | **<0.001** |
| High Frequency - 2cm | 0.03 | $4.81 \times 10^{-3}$ | 6.49 | **<0.001** |
| No Optic Flow - 2cm | -0.02 | $3.64 \times 10^{-3}$ | -6.69 | **<0.001** |
| High Frequency - Checker | 0.01 | $5.13 \times 10^{-3}$ | 2.82 | **0.005** |
| No Optic Flow - Checker | -0.04 | $4.06 \times 10^{-3}$ | -10.14 | **<0.001** |
| No Optic Flow - High Frequency | -0.06 | $4.42 \times 10^{-3}$ | -12.58 | **<0.001** |
| **Random effects** | **Variance** | **SD** | **R** | |
| Individual (intercept) | $2.90 \times 10^{-4}$ | 0.02 | | |
| Position(P2-P1) | $9.69 \times 10^{-5}$ | 0.01 | -0.43 | |
| Position (P3-P1) | $2.08 \times 10^{-4}$ | 0.01 | -0.39 | |
| Position (P3-P2) | $2.27 \times 10^{-4}$ | 0.02 | -0.34 | |
| Residuals | 0.25 | 0.49 | | |

Results from the linear mixed model fitted with Gamma family. Fish ID was added as random intercept and start position as random slope. Model selection on the random effect was conducted with ANOVAs (REML fitted) on pairs of models containing either “individual” as random intercept, “individual” and “start position” as crossed random intercept or “individual” as random intercept and “start position” as random slope. We used AIC criteria to select the best model .Significant results are in bold, *P* = *P* value; SD = standard deviation; SE = standard error; z = *z* value.

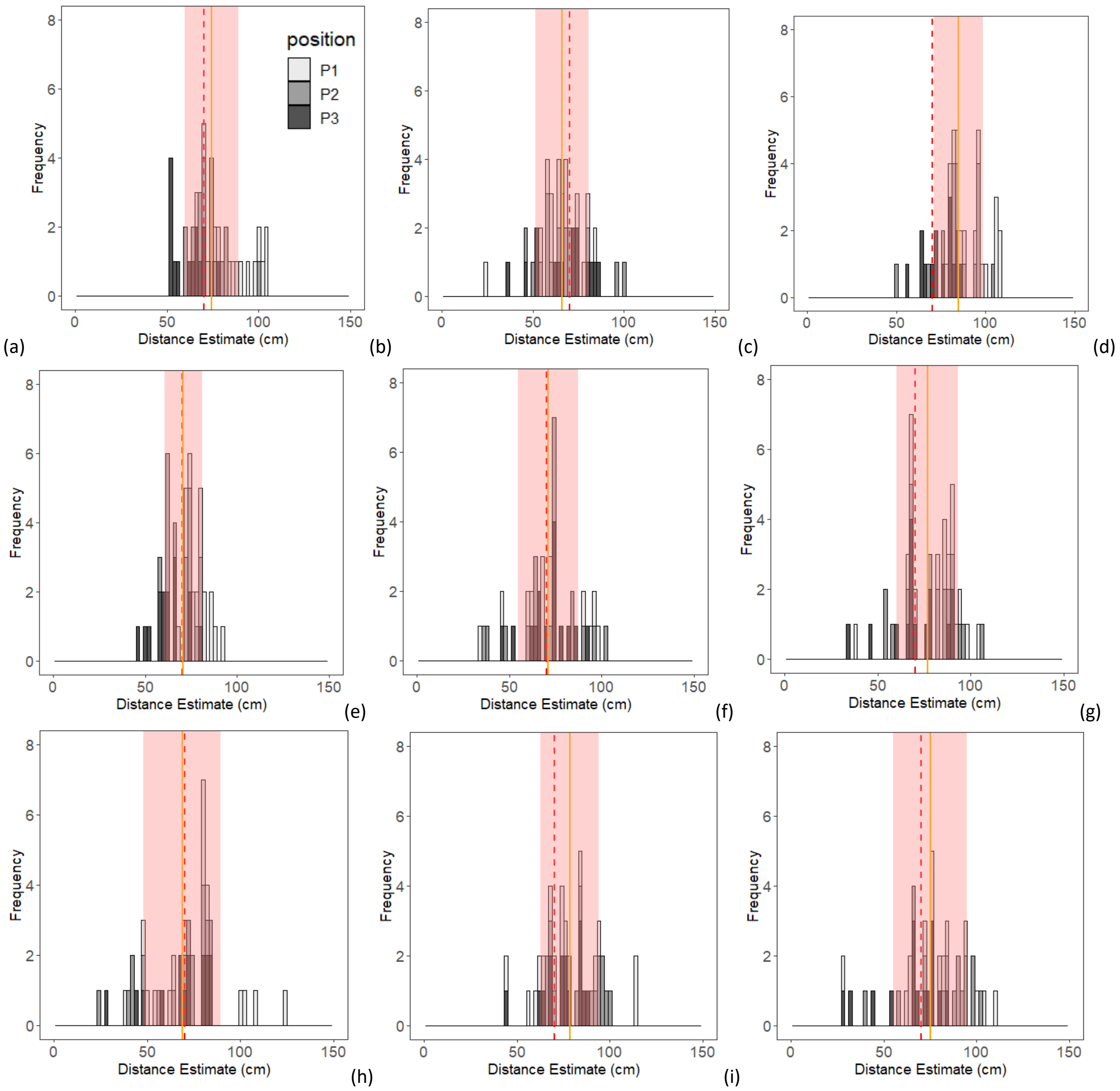


Figure A1: Goldfish distance estimation results with 2cm wide optic flow pattern. a,b,c,d,e,f,g,h,i, show results for G1,G2,G3,G4,G5,G5,G6,G7,G8 and G9 respectively. The red dashed line represents the target 70cm distance. The orange vertical line represents the mean fish distance and the pink shadow the SD around this mean. The shade of the bar indicates the start position of the test.

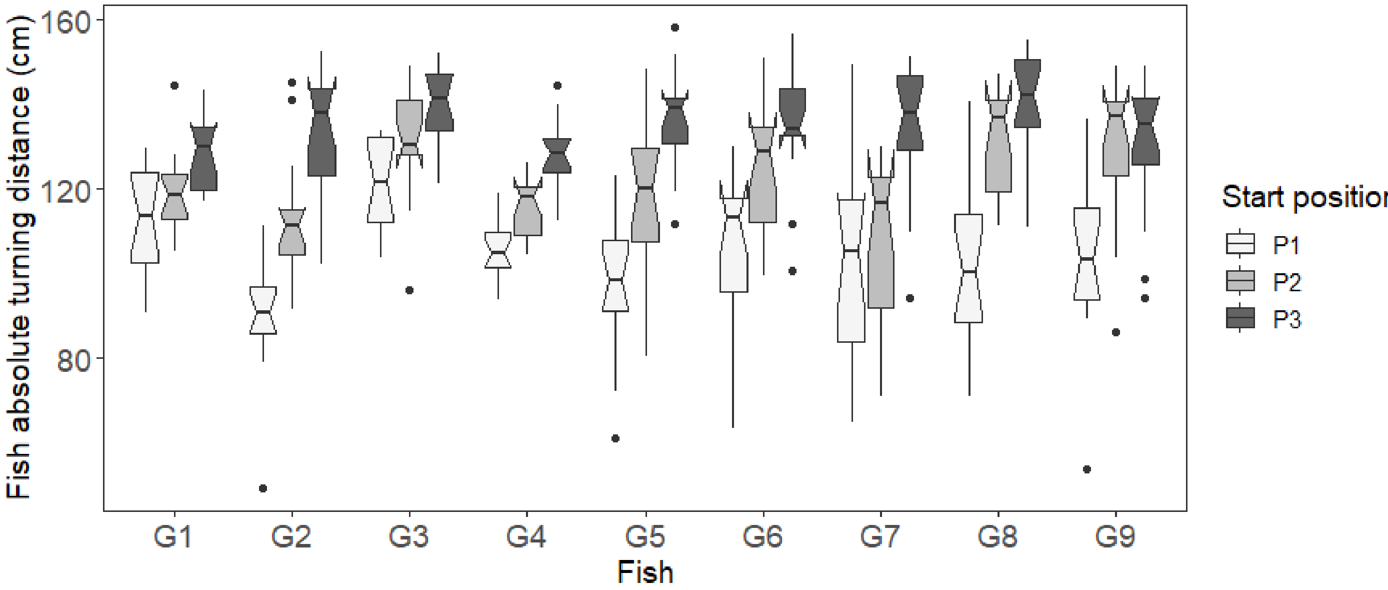


Figure A2: Goldfish absolute turning distance at each start position with 2cm Optic flow pattern. The difference in absolute turning position when individual start at the first (P1), second (P2) and third (P3) position indicate that the fish learned the target distance and are not using landmark cues. N=15 trials per position.

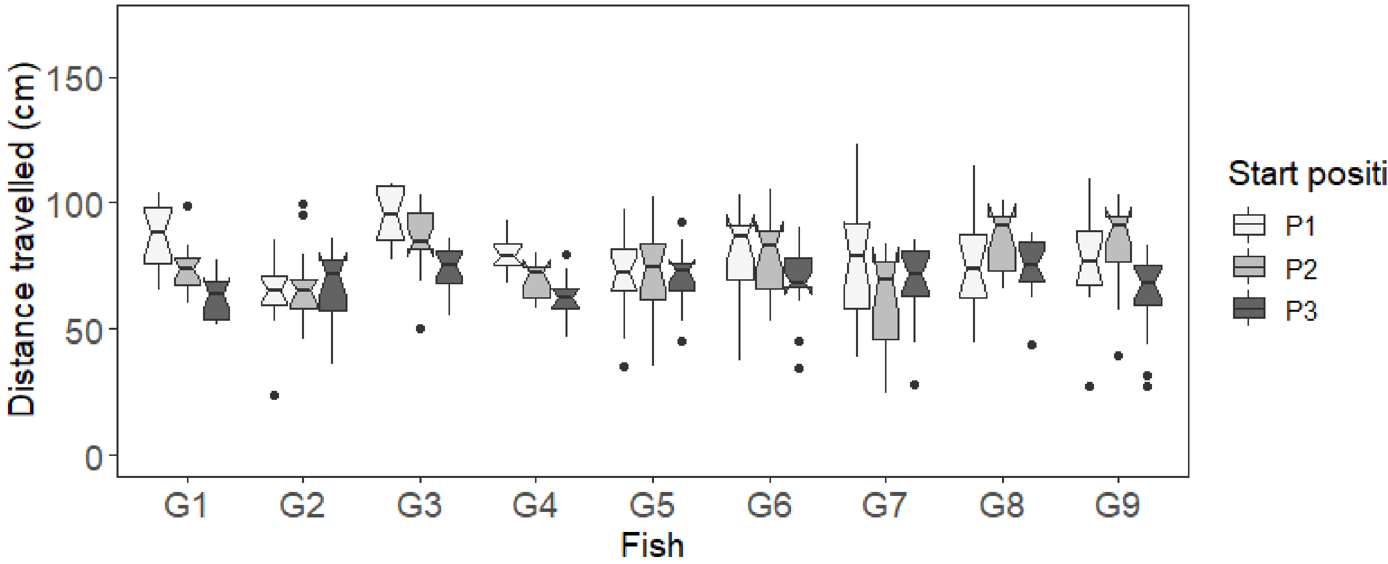


Figure A3: Distance estimation testing trials had three possible start positions to control for the possible use of external cues when solving the task. Distance travelled for each start position given the 2cm Optic flow pattern presented. N =15 trials per position.

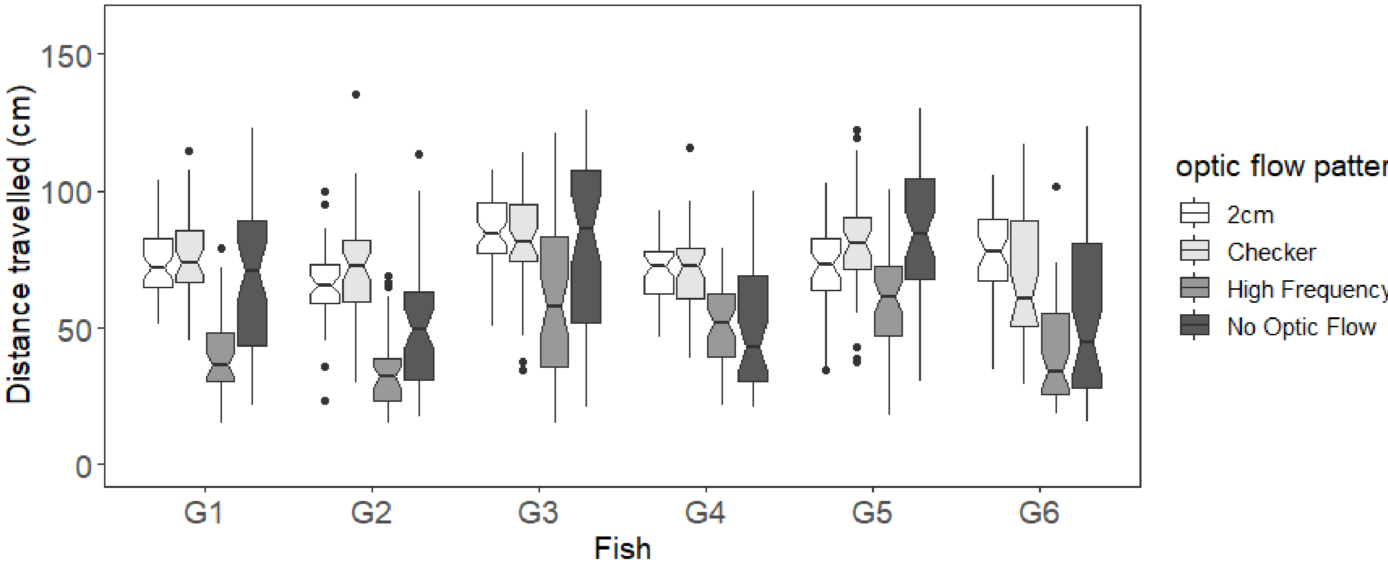


Figure A4: Distance estimate for the six fish tested with the different Optic flow patterns

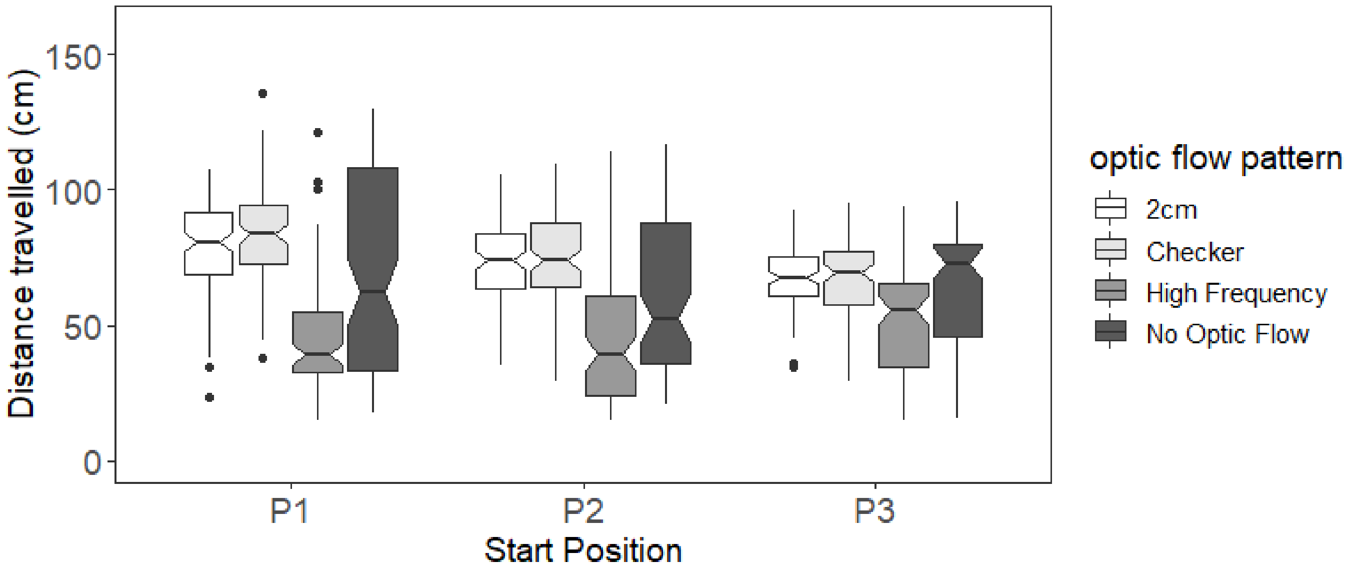


Figure A5: Distance estimate at the three start position with the different Optic flow patterns. N=6 individuals tested.
